# Noninvasive Stimulation of Peripheral Nerves using Temporally-Interfering Electrical Fields

**DOI:** 10.1101/2021.12.14.472557

**Authors:** Boris Botzanowski, Mary J. Donahue, Malin Silverå Ejneby, Alessandro L. Gallina, Ibrahima Ngom, Florian Missey, Emma Acerbo, Donghak Byun, Romain Carron, Antonino M. Cassarà, Esra Neufeld, Viktor Jirsa, Peder S. Olofsson, Eric Daniel Głowacki, Adam Williamson

**Author notes:** correspondence; /.

## Abstract

Electrical stimulation of peripheral nerves is a cornerstone of bioelectronic medicine. Effective ways to accomplish peripheral nerve stimulation noninvasively without surgically implanted devices is enabling for fundamental research and clinical translation. Here we demonstrate how relatively high frequency sine-wave carriers (3 kHz) emitted by two pairs of cutaneous electrodes can temporally interfere at deep peripheral nerve targets. The effective stimulation frequency is equal to the offset frequency (0.5 – 4 Hz) between the two carriers. We validate this principle of temporal interference nerve stimulation (TINS) *in vivo* using the murine sciatic nerve model. Effective actuation is delivered at significantly lower current amplitudes than standard transcutaneous electrical stimulation. Further, we demonstrate how flexible and conformable on-skin multielectrode arrays can facilitate precise alignment of TINS onto a nerve. Our method is simple, relying on repurposing of existing clinically-approved hardware. TINS opens the possibility of precise noninvasive stimulation with depth and efficiency previously impossible with transcutaneous techniques.

## 1. Introduction

Electrical stimulation of the nervous system is a powerful tool in fundamental biomedical research and is also at the center of bioelectronic medicine. Peripheral nerve stimulation (PNS) is at present one of the most extensive clinical fields of bioelectronic medicine, with the scope of new applications constantly growing^1–3^. Established examples of targets for PNS are the sacral, sciatic, and the vagus ^4,5^. Vagus nerve stimulation is clinically approved for treatment of certain types of epilepsy, and is in clinical trials for treatment of chronic inflammatory conditions, depression, arthritis, obesity, and other examples^6–11^. Sacral nerve stimulation has been utilized since the 1970s and is used to treat various bowel and bladder dysfunctions^12^. Foot-drop electrical therapy relies on stimulation of the nerves in the leg in order to treat foot drop, a condition where the feet “drag” and walking is difficult. Peripheral nerve stimulators are also used for treatment of various chronic pain disorders^13,14^. Presently, at both the level of fundamental research as well as clinical practice, PNS is relatively invasive. Implantation of stimulation electrodes, interconnects, and power supplies is necessary. Many of these techniques revolve around repurposing of well-established implanted cardiostimulator technology. Though these procedures are highly optimized and constantly improving^15,16^, surgery inevitably involves risk and patient discomfort. This is particularly problematic as many protocols require regular surgical battery changes, and additional invasive surgery if patients elect to have stimulators removed. For this reason, less invasive (minimalistic devices) or completely noninvasive stimulation solutions are of great interest. If the target nerve is not too deep below the skin, currents delivered from cutaneous electrodes can accomplish stimulation. This principle is behind transcutaneous electrical nerve stimulation (TENS)^1,14,17^. As a noninvasive solution, TENS procedures have been reported in numerous clinical studies, and commercial systems exist on the market. While very popular, the TENS approach has drawbacks. It is only applicable to shallow targets, and there are limits to the current magnitudes which can safely and comfortably be applied to the skin. Due to these limitations, as well as difficulty in precise targeting of nerves, widespread effective application of TENS has remained elusive, with the literature reporting mixed efficacy^18,19^. An alternative approach involves the use of focused ultrasonic waves, which can stimulate deeper targets than cutaneous electrical stimulation^20,21^. While promising, especially for central nervous system stimulation, for peripheral targets ultrasound has been less frequently used^22^. There is concern about ultimate safety of relatively high-power acoustic waves due to thermal and cavitation effects. Confounding effects from auditory stimulation via sound waves has also been reported as an obstacle. Alternative approaches based on transduction of near-infrared light are promising, however require specialized equipment and are in an early stage of development.^23–26^ The goal of effective noninvasive PNS remains an important unresolved issue in bioelectronic medicine.

In this work, we report a significantly more efficient transcutaneous electrical stimulation method – temporal interference nerve stimulation (TINS). This technique relies on two pairs of cutaneous electrodes driven by conventional clinical bipolar constant current stimulators. We exploit the principle of high-frequency temporally-interfering (TI) electric fields to stimulate deeper targets more efficiently than a typical topical nerve stimulator. TI is a relatively new noninvasive method, to-date explored in only a handful of brain stimulation experiments in animal models following its introduction by Grossman and coworkers in 2017^27,28^. TI stimulation relies on multiple high-frequency electric fields that only cause neuronal activation where they constructively overlap. By controlling field orientation and frequency offset, the hot-spot of constructive interference can be precisely targeted. The key aspect of this method is the use of carrier waves at frequencies higher than 1 kHz. Frequencies above this range are regarded as non-stimulating, and pass through tissues with relatively low loss. While these higher frequencies do not stimulate neural tissue, the interference envelope of two phase-shifted frequencies can elicit action potentials because the offset (aka “beat”) frequency can be tuned accordingly to < 100 Hz. These low-frequency envelopes stimulate neurons due to the nonlinear rectification of excitable cell membranes^29^.

Herein we test the TINS method on the mouse sciatic nerve, as this model allows rapid validation via observed muscle movement and electromyography (EMG) recording. TINS is applied via two pairs of surface electrodes driven by two clinical stimulators: one using a 3 kHz sine wave carrier frequency, and the other stimulator with frequency set to 3 kHz + *n* Hz offset (Figure 1A). We find that using the same electrodes and applying traditional TENS does not evoke a PNS response for the same amplitude of stimulation used with TINS. This corresponds well with finite element modeling of the electric field distribution, which demonstrates how TINS focuses electrical stimulation much deeper than TENS (Figure 1A). Having proved the efficacy of our approach using standard electrode pins, we next fabricated flexible multielectrode arrays (fMEAs). The fMEAs conform to the skin and have a grid of 40 addressable electrodes (Figure S1) which allow for selecting a combination of electrode pairs that optimally directs TINS to the peripheral nerve target (Figure 1B).

**Figure 1:**
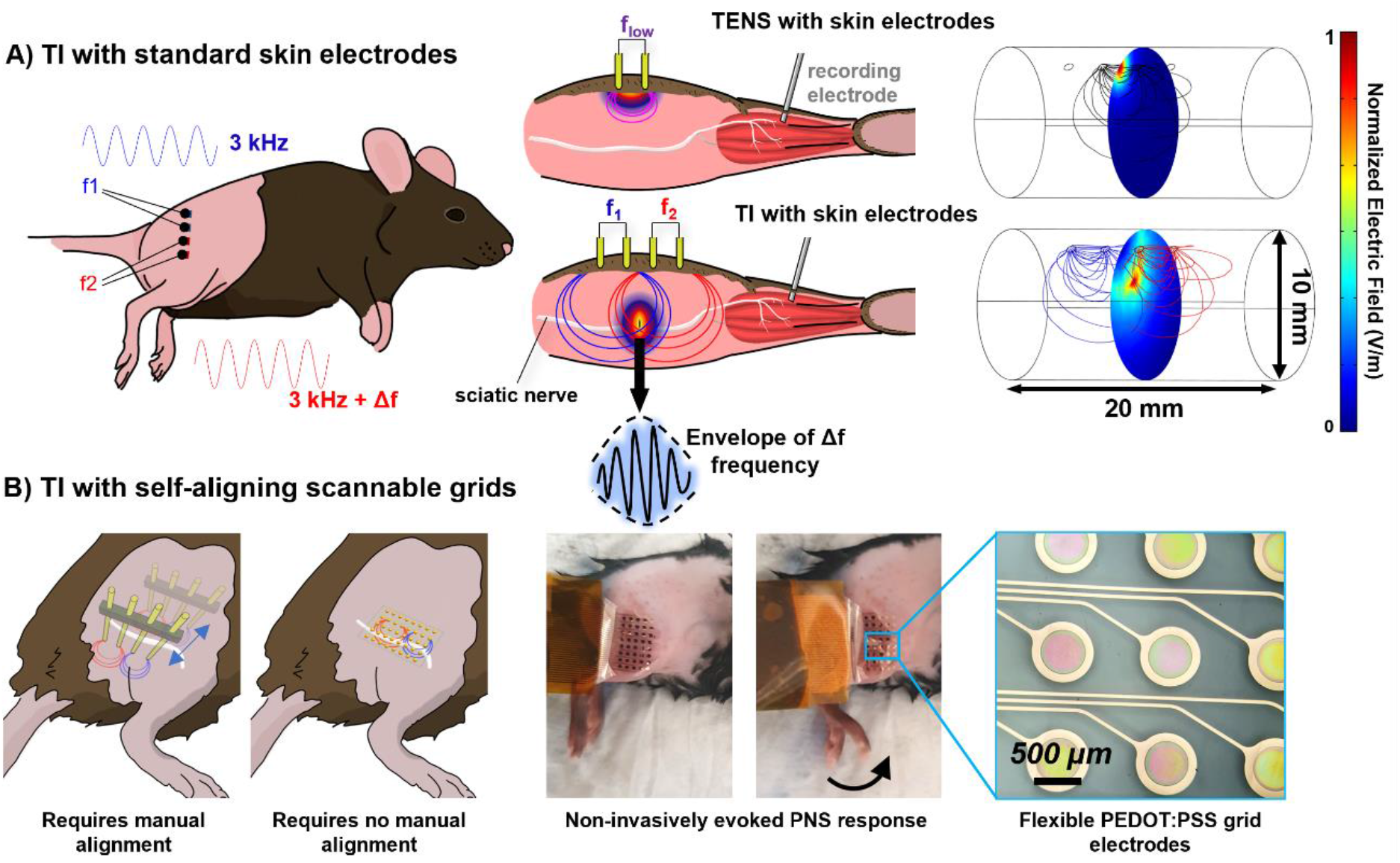
Temporal Interference Nerve Stimulation, TINS. The mouse sciatic nerve is used as a model PNS target, allowing both limb movement and EMG as read-outs. A) *Left:* TINS utilizes two pairs of electrodes driven at high frequency (3 kHz), with a frequency offset *n =* Δ*f*. The *n* becomes the interference envelope beat frequency which accomplishes stimulation at the hot-spot where the two electric fields interfere (*middle, bottom*). TINS allows reaching targets deeper below the skin than transcutaneous electrical nerve stimulation TENS (*middle, top*). *Right*: The calculated difference between TENS and TINS using finite element modeling of the electric field distribution on a cylindrical body. TENS generates the highest electric field in the immediate vicinity of the surface, while TINS with two electrode pairs results in a deeper focus of the electric field and the field near the surface of the cylinder is minimal. B) Stimulation electrode comparison: We first evoke PNS responses noninvasively using TINS from standard surface electrodes which are placed on the skin and positioned using a stereotaxic arm (leftmost image). To optimize alignment without needing any mechanical manipulation, we developed a flexible multielectrode array, fMEA, (4 μm-thick) with 40 conducting polymer coated electrodes which conform to the skin. *Right*: With the fMEA, optimal stimulation alignment can be accomplished using electronics alone.

## 2. Results and discussion

The first experimental goal was validating that TINS can work to stimulate the sciatic nerve. Experiments were performed using two clinically-approved bipolar constant-current stimulators (Digitimer DS5), modulating two separate pairs of electrodes. As stimulating electrodes, we initially utilized point electrodes in the form of gold pins (0.63 mm diameter), with a distance of 2.54 mm between each electode (center to center), as shown in Figure 1. Using a current of 350 μA with 3 kHz sine carriers and frequency offsets of 0.5 Hz – 4 Hz, we observed clear and regular muscle contractions and leg movements at intervals corresponding to the envelope frequency (Figure 2, and Supplementary Video S1). As a control, traditional TENS was attempted with two electrodes stimulating at 0.5 Hz – 4 Hz, and no PNS effect was observed using currents of 350 μA. Clear muscle movement was not evoked with TENS until >1 mA was applied.

**Figure 2:**
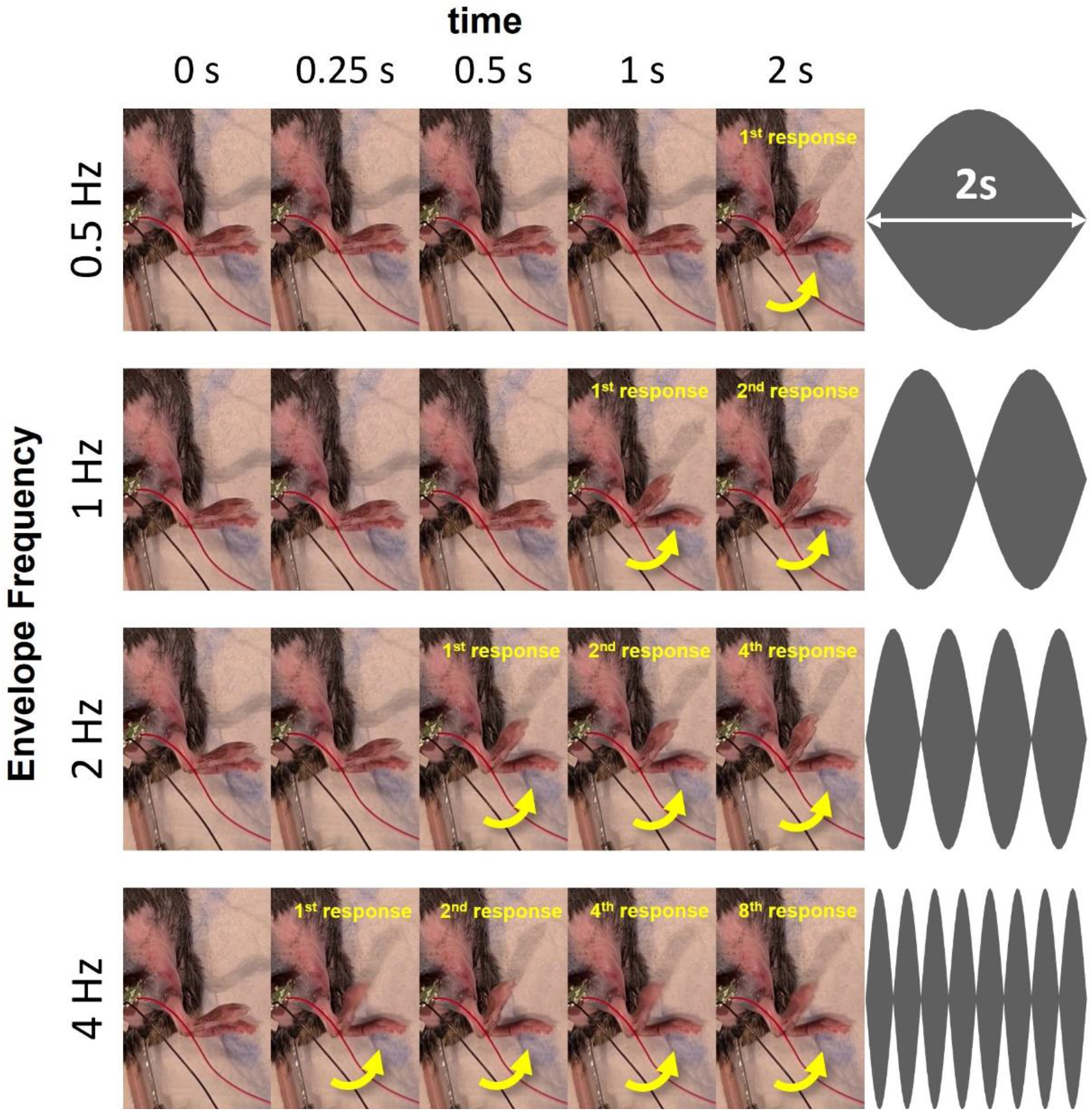
TINS evokes responses at the envelope frequency. Carrier frequencies of 3000 Hz and 3000 + *n* Hz (350 μA) are applied non-invasively to gold pin electrodes above the sciatic nerve. *n* is increased from 0.5 Hz to 4 Hz and the PNS kicking response follows the increase in envelope frequency. The video of this experiment is available in the supplementary information.

The TINS method was effective as shown in Figure 2, however the electrodes had to be carefully landmarked and aligned manually in order to overlap the interference hot-spot onto the sciatic nerve. To overcome the necessity of having to position the stimulation electrodes manually by manipulating a stereotactic arm, we fabricated a fMEA with a grid of 40 electrodes (5×8 electrodes, diameter 500 μm, spacing 1.25 mm between each electrode center to center), as shown in Figure 1B. The fMEAs are conformable grids that can be positioned roughly over the area where PNS is desired, and then pairs of electrodes can be cycled electronically to optimize targeting (Figure 3). The alignment of the stimulation was done by scanning TINS over different electrodes while recording EMG signals and monitoring motor movement with video. It should be noted that combinations of various electrodes allow the TINS to not only scan laterally to modify the *xy* position of stimulation hot-spot, but also vertically to reach deeper or shallower nerve targets. As seen in Figure 3A, alignment in this application utilizes video detection of the TINS evoked kick and a measured compound muscle action potentials (CMAPs). The effects of alignment can likewise be visualized in Figure 3A and by finite element modelling in Figure 4. The CMAP signal is the sum of numerous simultaneous action potentials from muscle fibers activated via a nerve, in this case the sciatic nerve. When an optimal alignment is detected, a CMAP is visible on the recorded signal or a vigorous kick is observed, the scan stops. Further parameters, for example the envelope frequency or stimulation amplitude, can then be studied on the nerve of interest. Using CMAP recording to tune TINS is enabled by the fact that the TINS stimulation artefacts are present at a significantly higher frequency range than the electrophysiological signal of interest. The power spectral density (PSD) only contains the two carrier frequencies and no low-frequency content, as it is the superimposed sum of two high-frequency oscillations (Figure 3B). Therefore, when performing electrophysiological recordings, it is trivial to apply a filter to the high-frequency stimulation artefacts, thus obtaining the biologically relevant signal.

**Figure 3:**
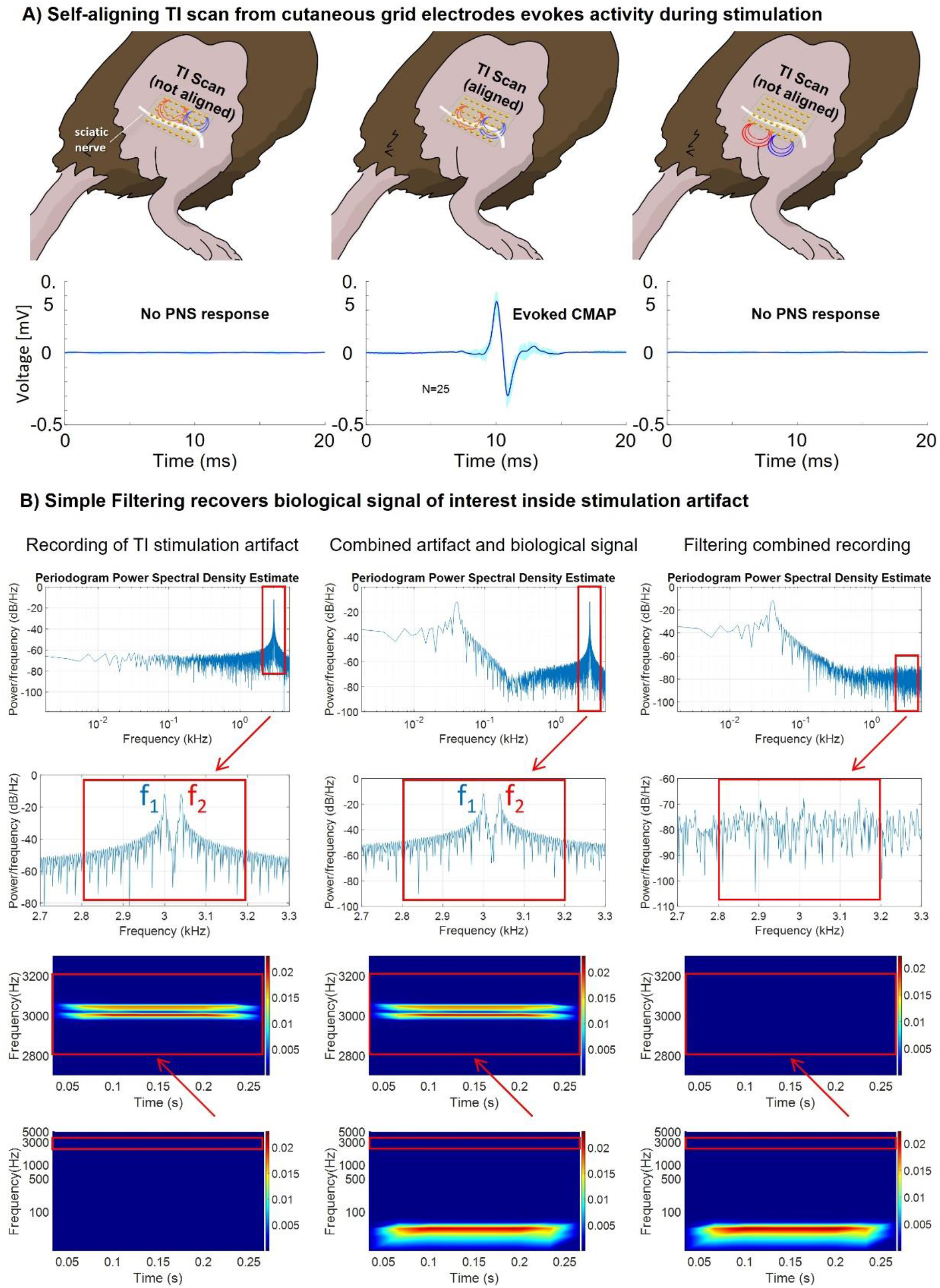
Self-alignment of TINS to deep peripheral nerves using scannable fMEAs. A) A Compound muscle action potential (CMAP) response is evoked and recorded synchronously with the leg kick. The scannable fMEA grid allows alignment to be automated, reducing the time required to locate the nerve. B) TINS creates electrical artifacts at much higher frequencies than the biological signal of interest. The raw electrical artifact of a TI signal contains only peaks at the carrier frequency, the envelope frequency does not appear in the PSD. (**Left panel)** The two pairs of electrodes on the mouse’s thigh therefore create only f1 + f2, not the envelope frequency and not other combinations of f1 and f2. Any other signals are thus from other sources, such as the electrophysiological activity of biological tissue **(Middle panel)** In PNS stimulation we selected sufficiently high carrier frequencies to be far away from the biological signal of interest. **(Right panel)** We see the resulting filtered PSD, leaving only the biological signal of interest. This phenomenon allows excellent recordings of events during stimulation and opens numerous opportunities for closed-loop phase-locking applications in PNS and prosthetics.

**Figure 4:**
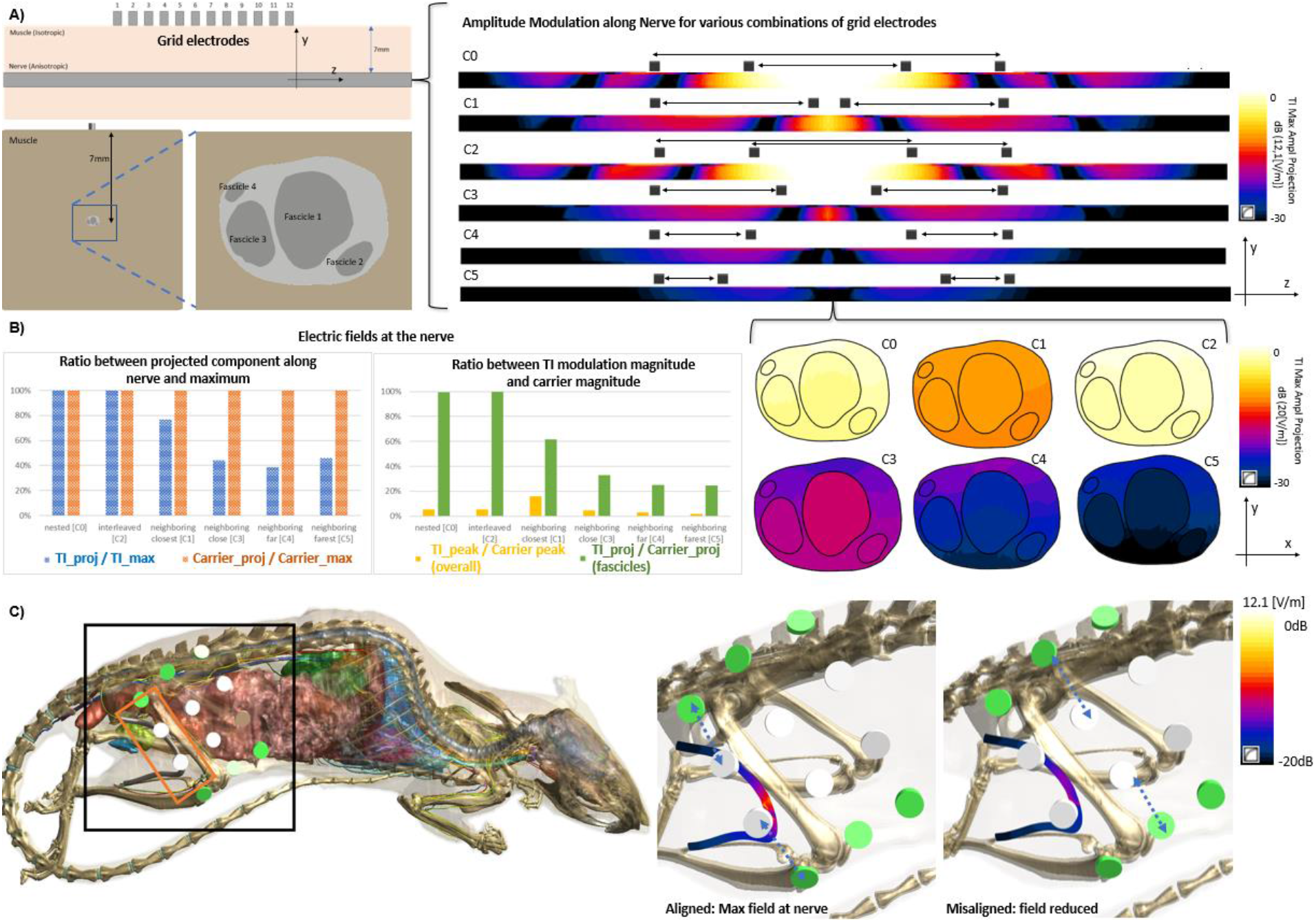
Simulation of Electrode Configurations and Electric Field at the Nerve. The selected configuration in from a row of electrodes, and the selected row parallel to the nerve of interest, strongly impacts the magnitude and profile of the amplitude modulation at the depth of the nerve. **A)** Schematic representation of the simplified simulation setup used to compare different stimulation strategies. It features a multi-fascicular nerve model at a depth of 7mm within muscle tissue and a range of electrode contacts. On the right, TI modulation envelope magnitude of the axon-aligned field component according to Equation 2 for different electrode configurations of the two electrode pairs (1 mA input current per pair; i.e. 2 mA in total with a current ratio of 1:1); nested (C0, electrode pairs 1-12, 4-8), adjacent (C1, electrode pairs 1-6, 7-12), interleaved (C2, electrode pairs 1-8, 4-12), adjacent with varying separation distances (C3, electrode pairs 1-5, 7-12; C4, electrode pairs 1-4, 8-12; C5 electrode pairs 1-3, 9-12). Electrode diameter 500 μm, **B)** left: ratio between the TI modulation amplitude (blue) of the field component aligned with the nerve fibers (Eq. 2) and the maximal one along any orientation (Eq. 1), as a measure for field alignment efficacy; orange: the same for the carrier field magnitude; right: ratio between peak TI modulation magnitude and peak carrier field magnitude (yellow: overall, green: within the fascicles), as a measure of TI vs. low-frequency stimulation efficacy at the target and outside. The nested and interleaved combinations provide the largest TI amplitude inside the nerve. **C)** Selection of a row of electrodes off-axis with respect to the nerve, as seen in the full simulation, rapidly reduces the TI-exposure of the nerve.

The TI field distribution for a range of electrode configurations was simulated to assess focality and nerve exposure amplitude assessed using computational modeling, comparing nested, interleaved, and adjacent (with varying distance) electrode pairs (Figure 4). Based on these calculations, it can be said that TI provides a much better balance between nerve stimulation and stimulation of overlaying tissues (by one order of magnitude). While a configuration with opposed electrode pairs could fully eliminate surface TI hot-spots, it was not suitable to to anatomical access limitation. All the other simulated configurations produce surface hot-spots where the fields from neighboring electrodes of different pairs are oriented parallelly. The TI magnitude at depth (according to Eq.s 1 and 2) is highest for the nested and the interleaved configurations (which are also the least focal ones). For these configurations, the TI magnitude is comparable to the carrier magnitude (again because of the parallelity), while it drops to 20% for the most separated variant of the adjacent configurations. At the same time, the TI modulation magnitude at the surface can be more than one order of magnitude smaller than that of the carrier field. The adjacent one with the smallest separation distance is the most focal one, but also produces the strongest surface TI exposure; surface exposure quickly diminishes with increasing separation, while focality is maintained. When considering the strong preferential axon orientation in nerves and evaluating the TI modulation amplitude exclusively along the nerve direction, the efficiency of the the adjacent configurations is strongly reduced (up to a factor >2) compared to the nested and interleaved ones, where currents are better aligned with the nerve in their intersection region. Figure 4 also illustrates the impact of the fascicular nerve structure, with its semi-insulating epineurium on exposure strength. The perineurium, combined with the anisotropic fascicular conductivity, result in a longitudinal smoothing, but also an overall reduction, of the intrafascicular fields. In summary, when it is not possible to place the second electrode pair on the opposing side for anatomical reasons (too distant), the nested configuration is preferable in terms of stimulation effectivity and surface exposure reduction when focality does not matter, while the adjacent configurations (at an optimized separation distance) are preferable otherwise.

## 3. Conclusions

The tradeoff of non-invasiveness versus specificity is an accepted limitation in the field of electrical stimulation of the nervous system. Highly selective stimulation at anatomic depth usually requires invasive, implanted electrodes^5^. Non-invasive treatments, meanwhile, necessarily sacrifice specificity. The main motivation of our work has been overcoming this fundamental obstacle. The TINS method we have tested, which relies on safe transmission of high frequency signals through the tissue and subsequent constructive interference at the stimulation area of interest, can indeed solve this problem. Our data validates efficacy of stimulation of the sciatic nerve, under conditions where normal transcutaneous stimulation, TENS, could not elicit any response. The TINS method is supported by modeling and calculation, and relies on relatively well-established principles in electrodynamics. Furthermore, TINS can be performed using existing hardware, provided proper channel isolation and very good linearity are ensured, which makes adoption in animal research and potential clinical translation facile. The geometry of the electrode pairs used in TINS dictates where the stimulation hotspot will be. To facilitate precise nerve targeting, we have demonstrated the use of soft and flexible multielectrode arrays which can be roughly positioned over the area of interest. Scanning of different permutations of electrode pairs can then be done automatically, while performing read-out of desired stimulation effects. This procedure is made more powerful by the fact that the stimulation artefacts produced by TINS are exclusively in a high-frequency range ≥ (3kHz), allowing filtering to be used to isolate electrophysiological signals of interest. TINS allows non-invasive stimulation experiments to be performed using experimental animal models, thus enabling large-scale neuromodulation experiments that were previously extremely laborious or impossible. We envision TINS as a useful clinical procedure for benchmarking the efficacy of electrical stimulation in patients who are candidates for an implantable technology. TINS can allow clinicians to test and titrate stimulation parameters to determine if patients are good candidates for a more invasive therapeutic bioelectronic intervention. Finally, besides these examples of acute applications, advances in technology of conformable cutaneous electrodes^30,31^ can allow TINS to be applied chronically in bioelectronic medicine applications. Future research to expand the possibilities with TINS can feature using more than two electrode pairs to further focus the stimulation hot-spot.

## 4. Experimental Methods

### Animal Procedures

This study and all experimental protocols were approved by the *Stockholm Regional Board for Animal Ethics* (Stockholm, Sweden). Three mice were used in these experiments.

### Fabrication of flexible multielectrode arrays

The flexible, ‘self-aligning’ electrode grid fabrication process is based on previously reported methods.^32,33^ First, a parylene-C (PaC) film was deposited on a clean glass wafers using a chemical vapor deposition system (Diener GmbH), resulting in a thickness of 2.5 μm. No adhesion promoter is used as this PaC layer is delaminated from the glass after all fabrication steps are complete and acts as the final underlying substrate. Photolithography and lift-off processes were employed to pattern metal interconnects on top of the PaC substrate. This was performed using a negative lift-off photoresist (nLof 2070), a SUSS MA 6 UV broadband contact aligner and AZ MIF 729 developer. After development, an O_2_ plasma treatment is carried out and 2 nm chromium adhesion promoting layer and 100 nm of gold were then thermally evaporated onto the substrates. The samples were immersed in an acetone bath to complete the lift-off process and define the interconnects. To electrically insulate the metal lines, a second parylene-C (PaC) film was deposited on the devices to thicknesses of 2 μm, using the same deposition process as before, however with 3-(Trimethoxysilyl)propylmethacrylate present in the chamber to act as an adhesion promoter. Subsequently, an etch mask is patterned using the positive photoresist AZ 10XT patterned with the UV contact aligner and AZ developer. This is used to define the outline/shape of the overall device and the PaC is etched through reactive ion etching (RIE) with an O_2_/CF_4_ plasma. Following this etch and subsequent removal of the photoresist, a dilute solution of Micro-90 industrial cleaner was spin coated onto the insulation layer, followed by the deposition of a sacrificial PaC layer (2 μm). These steps allow the sacrificial layer to later be peeled from the substrate, defining PEDOT:PSS at the electrode sites. The sacrificial PaC layer was etched as before, using RIE after an AZ 10XT mask was patterned to open the electrode sites and back contacts by creating an opening down to the gold interconnects. A dispersion of PEDOT:PSS (CleviosTM PH 1000 from Heraeus Holding GmbH), 5 volume % ethylene glycol, 0.1 volume % dodecyl benzene sulfonic acid, and 1 wt % of (3-glycidyloxypropyl)-trimethoxysilane was spin coated onto the substrates. Multiple layers were coated to attain a thickness of ∼1.5 μm allowing good skin contact of the final device without the use of electrolyte. Between each coating process, short baking steps were used (1 min, 90 C) and the substrates were allowed to cool to room temperature. The sacrificial PaC layer was peeled off removing superfluous PEDOT:PSS and defining the electrode sites. The devices were baked for 1 h at 140 C to crosslink the film. Finally, the devices were immersed in deionized water to remove low molecular weight compounds. This water soak also facilitates the delamination of the final flexible electrode array from the glass wafer.

To make electrical contact to the flexible devices, the backside was laminated onto a Kapton film. This was placed into a zero insertion force clip (ZIF-Clip®) soldered onto an adapter printed circuit board. This board allowed for simple wired connection to current source.

### Recording of CMAPs in the mouse leg

Measurements were obtained using a 27G stainless steel recording needle implanted in the *tibialis anterior* muscle and a reference needle implanted in the skin above the extensor *digitorum longus* muscle. Both needle shafts were passivated with insulating plastic, with only 2 mm remained uninsulated at the tip. Both electrodes were connected to an RHS Stim/Recording System from *Intan technologies* through a 16-channel RHS headstage. Recordings were captured with a 50 Hz notch filter.

### Mouse Preparation

Mice were anesthetized with isoflurane gas, a concentration of 3% was used to induce anesthesia and a concentration around 1.5 - 2% was used to maintain the anethesized condition. Mice were then placed slightly on the side to allow a comfortable access to the leg. The mouse position was secured utilizing cushions of gauze under their belly and neck. The mouse temperature was monitored at 37°C and maintained through a Homeothermic System from *AgnThos*. Hair Removal Cream from *Veet* was used to remove fur on the leg and then gently rinsed with water.

### Temporal interference electrical stimulation, TINS, of the sciatic nerve

Temporal interference stimulation of sciatic nerve was done using a 4-pin gold-plated contacts header from Farnell (ref. 825433-4), each of the contacts is separated by a center-to-center distance of 2.54 mm. The pin header was placed above the sciatic nerve with the 4 contacts touching the skin. Special care was made to ensure that the pin header was aligned with the sciatic nerve, when the position of the header seemed to be ideal, it was maintained thanks to a third hand soldering helper. If stimulation turned out to be ineffective, the pin header was repositioned until the stimulation drove a muscular response. Stimulation parameters were provided by a two-part system: Waveforms (frequency and waveshape components) were provided by a function generator *Keysight* EDU33212A and current amplitude was controlled by a DS5 isolated bipolar constant current stimulator from *Digitimer*. Each pair (the stimulator and its ground) of electrode was connected to its own DS5, itself connected to its independent function generator.

TINS of the sciatic nerve was also done using a conforming grid of conducting polymer electrodes. Once the grid was placed on the leg mouse it was possible to stimulate through any of the 40 contacts composing the grid. This allowed the manipulator to dynamically select different spatial configuration and eliminated the need to reposition the grid if stimulations were ineffective. Stimulation parameters were provided in the same fashion as with the pin header.

### Signal treatment

Signals were recorded in .rhs format and converted in .mat then analyzed with MATLAB (Mathworks) R2020B. Signals were lowpass-filtered at 800 Hz to remove the high-frequency stimulation artefact. All signals were aligned to the positive maximum of their CMAP before any statistical calculus. 25 typical CMAP have been extracted and the average response has been plotted along its calculated interquartile area.

### Finite element modeling

Finite element model simulations were done with COMSOL Multiphysics 5.5 for Figure 1 and Sim4life v6.2.1.4972 (ZMT Zurich MedTech AG, Zurich, Switzerland) for Figure 4. Both, simplified (more controlled) and anatomically detailed (more realistic) modeling was performed. The former features a 2.5 dimensional nerve model, generated by linearly extruding a manually segmented histological cross section image of a rat sciatic nerve, embedded at 7mm depth in a homogeneous volume conductor filled with muscle tissue (σ_axial_=0.33 S/m). The latter involves the anatomically detailed“Neurorat” rodent model (obtained by segmenting MRI and CT image data from a 150g and 150mm long (without tail) Dark Aouti rat^34^ in which a nerve model was embedded by extruding the same cross section used in the simplified model along the model-provided sciatic nerve trajectory. Dielectric properties were assigned according to the low-frequency dielectric tissue parameters from the IT’IS tissue properties database^35^ and the predefined tissue assignment tags in the NEUROFAUNA model. Nerve properties were treated as anisotropic within the fascicles (i.e., differing longitudinal and transversal conductivities; σ_radial_=0.087 S/m and σ_axial_=0.57 S/m). Epineurium and fascicle perineurium were treated as thin, highly-resistive layers, with σ_memb_=0.00087 S/m and thicknesses assigned as 3% of the fascicle/nerve diameter according to ref. 36. Electric fields for the electrode pairs of interest were simulated using the ohmic-current dominated electroquasistatic solver family in Sim4Life, which solves the equation ∇σ∇ϕ = 0 – a valid approximation of Maxwell’s equation^37^ when the wavelength is large compared to the simulation domain and displacement currents are negligible compared to resistive ones (these conditions were verified for the given setups). Electrodes were modeled as cylinders sized and placed in accordance to the experiments and to the investigated scenarios from Figure 4. Dirichlet boundary conditions were used before normalizing the input currents to 1mA per pair (resulting in a total input current of 2mA), as they provide superior predictions of current distributions near electrodes compared to flux boundary conditions. Convergence analyses were performed to ensure that the resolution and solver tolerances are suitable. The built in Sim4Life functionality was used to compute the TI exposure metric according to [1]:

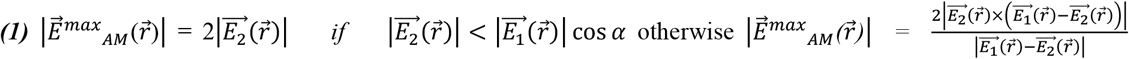

which is the maximum of the modulation envelope magnitude along any spatial orientation (assuming without loss of generality that *α* < *π*/2 and 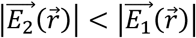). When there is a dominant neural orientation (e.g., axons in nerve), the modulation magnitude of the field component along that direction is used as TI exposure metric instead:

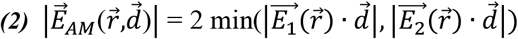

Simulations on the anatomical model were executed using the structured variant of the ohmic current dominated low-frequency solver (∼110 Miovoxs, 0.08-0.25 mm resolution). The simulations on the simplified model were performed using the unstructured solver variant of the low frequency (40 Mio tetrahedral, pyramidal, and prismatic elements), which is numerically less robust, but provides the thin resistive layer model used to model the perineurium and the epineurium.

For the COMSOL simulation in Figure 1, a cylinder mesh of 10 mm diameter and 20 mm of length was designed to represent a portion of a mouse leg. Each electrode interface with the mouse was defined by a disk of 0.63 mm. Dieletric properties for muscle tissue at 3 kHz, permittivity of 9.79E+4 and electrical conductivity of 3.33E-1 S/m were applied to the grid model. The physics interface choosen for the simulation was Electrical Currents Interface solver with this equation : *∇*.*J* = *Q*_*j*.*v*_, where 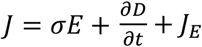 and *E* = −*∇V*. The study node selected was Time Dependent. All plots were directly obtained via the built-in tools of COMSOL Multiphysics 5.5.

## Acknowledgments

This project has received funding from the European Research Council (ERC) under the European Union’s Horizon 2020 research and innovation program (A.W. grant agreement No. 716867; E.D.G. grant agreement No. 949191). E.D.G. acknowledges funding from the city council of Brno, Czech Republic. M.J.D acknowledges funding from the European Research Council (834677 “e-NeuroPharma” ERC-2018-ADG). PSO and ALG were funded by MedTechLabs and the Swedish Reseaach Council.

## Author contributions

E.D.G. and A.W. conceived the project. B.B., and M.J.D performed experiments. M.J.D and D.B. designed and fabricated flexible cutaneous multielectrode arrays. B.B.; M.J.D.; A.L.G.; M.S.E.; F.M.; and E.A. all performed *in vivo* stimulation and recording experiments and analyzed data. B.B., I.N., A.M.C and E.N did the finite-element simulations and analysis. A.W.; E.D.G.; M.J.D and B.B., wrote the paper with input from all other authors. P.S.O.; E.D.G.; and A.W supervised the project.

## Supplementary figures

**Supplementary Figure 1:**
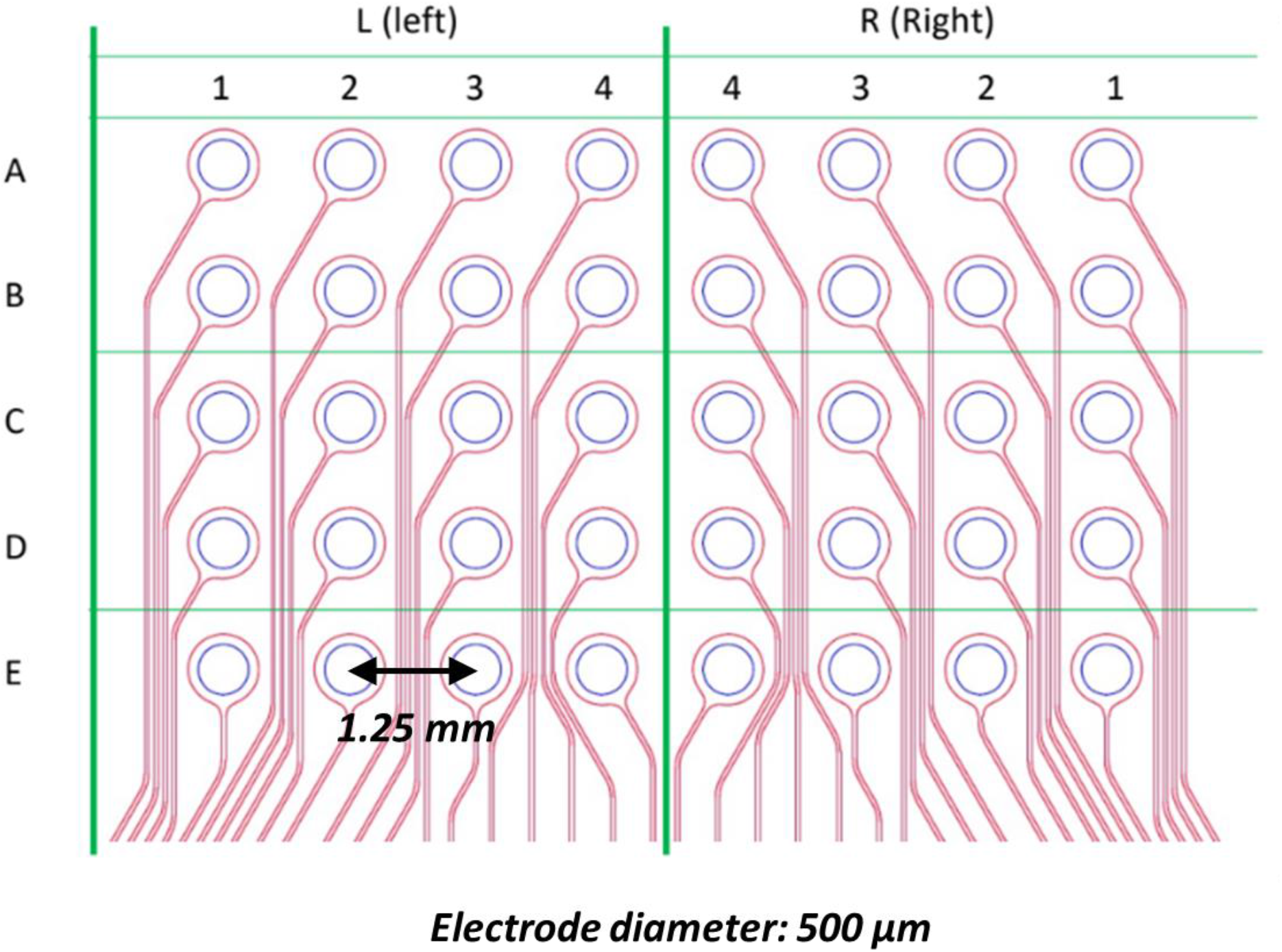
Spatial organization of the grid. Each contact can be configured to act as a current source or as a ground.

